# Modelling Microtube Driven Invasion of Glioma

**DOI:** 10.1101/2023.09.05.556421

**Authors:** Thomas Hillen, Nadia Loy, Kevin J. Painter, Ryan Thiessen

## Abstract

Malignant gliomas are notoriously invasive, a major impediment against their successful treatment. This invasive growth has motivated the use of predictive partial differential equation models, formulated at varying levels of detail, and including (i) “proliferation-infiltration” models, (ii) “go-or-grow” models, and (iii) anisotropic diffusion models. Often, these models use macroscopic observations of a diffuse tumour interface to motivate a phenomenological description of invasion, rather than performing a detailed and mechanistic modelling of glioma cell invasion processes. Here we close this gap. Based on experiments that support an important role played by long cellular protrusions, termed tumour microtubes, we formulate a new model for microtube-driven glioma invasion. In particular, we model a population of tumour cells that extend tissue-infiltrating microtubes. Mitosis leads to new nuclei that migrate along the microtubes and settle elsewhere. A combination of steady state analysis and numerical simulation is employed to show that the model can predict an expanding tumour, with travelling wave solutions led by microtube dynamics. A sequence of scaling arguments allows us reduce the detailed model into simpler formulations, including models falling into each of the general classes (i), (ii), and (iii) above. This analysis allows us to clearly identify the assumptions under which these various models can be *a posteriori* justified in the context of microtube-driven glioma invasion. Numerical simulations are used to compare the various model classes and we discuss their advantages and disadvantages.

## 1 Introduction

Glioblastomas constitute aggressive and fatal brain cancers, with the majority of patients dying within a year and a half of diagnosis and only a few percent surviving five years or longer [30, 49]. Standard treatment has remained largely the same for more than a decade, consisting of maximum safe resection of the (visible) tumour mass, radiotherapy, and chemotherapy [48, 49]. A major barrier lies in the highly invasive nature of gliomas, highlighted by a diffuse border and disease re-occurrence at or near the margins of the treated area [21]. Consequently, significant research has been devoted to the mechanisms leading to glioma cell dissemination [9, 20]. Glioma cells invade locally (migrating through the extracellular matrix, ECM), but also exploit oriented structures (e.g. vasculature, white matter fibre tracts) that facilitate longer movements [9]. As a consequence, glioblastomas typically present a heterogeneous and anisotropic shape that confounds precise determination of tumour extent.

Recently, a spotlight has been cast on the capacity of various cell populations to extend long thin nanotube structures (variously named tumour microtubes, tunneling nanotubes, cytonemes, membrane bridges) that allow direct connection and signalling between cells [37]. Within gliomas, certain astrocytomas form these tumour microtubes (TMTs), with their number and length (up to 500 *µm*) increasing with the extent of tumour progression [29]. The actin-rich and highly dynamic tubes infiltrate healthy brain tissue at the invasive front, with a significant fraction following oriented axons. Further, they aid tumour dissemination: divided nuclei travel down microtubes, settling at a location removed from the point of mitosis. Over time a network is formed, with previously unconnected cells integrated through coupling to microtube ends. The network matures to the point at which signalling occurs, long range cell to cell communication indicated through the observation of intracellular calcium waves. Notably, the emergence of this TMT-connected network in glioma bears resemblance to the networks formed during brain development and possesses a degree of self-repairing capacity, with greater resistance against both radiotherapy [29] and chemotherapy [47] treatments.

Elements of glioma growth and treatment have been the focus of numerous mathematical models over recent years, for example see the reviews [1, 20]. Given the phenomenal complexity of glioma growth – interactions between glioma cells and their microenvironment, immune cells, vasculature, molecular signalling systems etc, all embedded within a complex heterogeneous brain tissue – an all-encompassing model is a futile endeavour at present and models tend to consider well defined sub-processes [20]. Macroscopic partial-differential equation (PDE) models that predict invasive spread have formed a significant part of this literature, stemming from the work of Murray, Swanson and others (see the reviews of [43, 1]). Much of this work is founded on the highly influential PI-model (proliferation-infiltration model), a variation on the classical Fisher-KPP equation [27]. The key strength of this model lies in its simplicity, calling for a minimal set of parameters that facilitates its fitting to patient-specific data. Of course, the flip-side is lack of detail and various further models have been proposed.

One extension has been to model at the mesoscale, i.e. the scale of movements performed by individual glioma cells [33]. This approach allows preferred cell orientations to be incorporated, for example according to aligned neural axons or capillaries [9], and scaling yields a “fully-anisotropic” version of the PI model at the macroscale. When applied to environments informed from diffusion tensor imaging (DTI) datasets, this can lead to improved fit against patient-acquired imaging data [42]. Other anisotropic diffusion models have been formulated, for example motivated from classical phenomenological arguments (e.g. [23]) or extended to include further mesoscale detail (e.g. [11]).

An extension of these models is to incorporate the long-standing “go-or-grow” hypothesis for glioma growth, a proposed dichotomy between proliferation and invasion [14]. While experimental tests into the relevance of go-or-grow remain ambiguous (e.g. [14, 8, 18, 46]), models that incorporate this concept have been developed in [15, 34, 38, 40] amongst others, via splitting the glioma cell population into two states: actively proliferating, or actively migrating.

Another significant branch of modelling has been to explore the mechanical consequences of glioma growth, e.g. [4, 24, 19, 5, 41, 35]. While initial models used a linear stress-strain relationship, more recent models include nonlinear elasticity models such as the one-term Ogden model [35]. However, while mechanical effects are undeniably important, here we will focus our efforts at the diffuse leading invasion edge of a growing tumour. In this region, glioma cell densities are relatively low and mechanical effects are minimal. Mechanical effects become important once a bulk tumour of sufficient size is established [4, 24].

Implicit to the models described above is an assumption of independent invaders – migrating cells infiltrate at the front as individuals – and the resulting models possess a diffusive-type structure that resembles the non-compact border of gliomas. The observations of [29], rather, imply that there can be an interconnectedness to the invasion process (at least, for certain tumours): new tumour mass can arise from nuclei that migrate along the microtubes that extend from existing cells. Here we build a mesoscale model to describe the intricacy of this process. Specifically, our *glioma-TMT transport model* includes variables for the mature TMTs, the tips of extending TMTs, the bulk glioma cells, the migrating nuclei and the resting nuclei. We note that the explicit variables to describe TMT dynamics distinguishes our model from a recent approach in [6], where a flux-saturated model was proposed that accounts for the role of TMT protrusions and integrin-mediated ECM degradation at the invading front. Steady state and numerical analysis of the glioma-TMT transport model reveal its capacity to recapitulate features of glioma-microtube growth. In particular, we observe the formation of a travelling wave profile in which TMT tips infiltrate at the invasive front, which in turn leads to movement of nuclei along TMTs into this region and subsequent increase in the tumour bulk.

The complexity of this model, however, precludes an extensive formal travelling wave analysis. Consequently, we adopt a sequence of assumptions that allow reduction to simpler forms. First, we reduce to a simpler model for just the TMT tips and bulk population, but retain the mesoscale (the tips-bulk transport model). Second, we perform a parabolic scaling analysis that leads to a macroscopic reaction-diffusion model for the tip and bulk populations, which is either of anisotropic or isotropic diffusion form (the tips-bulk anisotropic or isotropic model, respectively). Finally, we reduce to a single reaction-diffusion model for just the bulk population, which can again be of anisotropic or isotropic form (the bulk anisotropic or bulk isotropic model, respectively). A notable consequence of the assumptions is that the resulting models fall into classes related to various previous models for glioma invasion: the tips-bulk models each have a go-or-grow type structure, the tips-bulk anisotropic and bulk anisotropic models have anisotropic diffusion structure, and the bulk isotropic model is of classical PI structure. In Figure 2, a pathway is laid out that passes through various models for glioma invasion, where successive models are obtained through specific assumptions regarding the biological processes included/excluded.

## 2 Glioma-TMT transport model

### 2.1 Observations for model formulation

Here we describe some key observations of TMT dynamics based on the study of [29], focusing on those most relevant to the model formulation. These observations are principally derived from a murine model of glioma development, accompanied by histological analysis of human tissue sections. Note that the data in [29] is accompanied by four supplemental videos, which were instrumental in our parameter estimation.

(O1) TMTs are primarily a phenomenon of gliomas of astrocytoma origin: for example, while a frequent feature of astrocytomas, they are infrequent within oligodendrogliomas. Further, they become increasingly present as the tumour progresses, with tissue sections of glioblastoma multiforme (GBM) indicating their presence in 93% of cases.

(O2) At the invasion front, TMTs infiltrate brain tissue in a highly dynamic and searching fashion, rapidly extending and retracting.

(O3) Both the number and length of TMTs increase with tumour progression, frequently with more than 4 per cell and TMTs occasionally extending more than 500 *µ*m from their point of origin. The actin-rich nature of TMTs indicates their potent motility, and thin TMTs frequently branch from mature ones.

(O4) TMTs are frequently used as tracks for the movement of nuclei, where mitotic events are followed by the separation of nuclei and movement along TMTs (with mean speeds 66*±*35 *µ*m day^*−*1^), potentially settling more than 100 *µ*m apart and becoming part of the bulk.

(O5) A significant fraction of TMTs follow axons, tightly coupling the structure of the TM network to inherent brain anisotropy.

(O6) With time, an increasing number of TMTs were found to sprout from one cell and connect to a second. This growing intercellular connection stems partly from the division and subsequent separation of nuclei inside the network, described above, but also from the pairing of open TMT ends to previously unconnected cells (anastomosis).

(O7) Increasing intercellularity opens the pathway to growing communication within the network, exemplified by long range calcium signalling waves. Notably, TMT networks appear to be more resilient against treatment. Damaged parts are identified within the network structure and a targeted healing process is triggered.

### 2.2 The full glioma-TMT transport model

We first remark that the formation of the TMT network bears resemblance to other processes of network growth in biology, ranging from hyphae growth in filamentous fungi (the formation of long, branched filament structures, that extend through the soil and increase nutrient uptake [39]) to angiogenesis, i.e. formation of new vasculature. Many continuous models for such processes rely on the “snail-trail” approach, developed in [10] to describe filamentous fungal growth, and subsequently adapted to model angiogenesis in [2]. Specifically, this model describes the network via the densities of two key variables: the “tips” of the network-forming structures, a motile population that moves through the environment, and the “stalks”, a sessile population laid down in the rear of tip movement. The name snail-trail refers to the similarity to the slime-trails left by snails, with the microtube tips representing the snail and the microtube representing the stalks.

Accordingly, we develop a kinetic-transport model for TMT driven growth, in which the network is represented by two variables: the TMT tips, *P* (*t, x, v*), and the (mature) TMTs, *M* (*t, x, v*), formed as the tip extends. In addition, we consider variables for migrating nuclei, *N* (*t, x, v*), the bulk tumour mass, *B*(*t, x*), and a pre-bulk state, *R*(*t, x*), during which nuclei have stopped migrating and mature into bulk. A schematic is given in Figure 1. In the above, each of *P* (*t, x, v*), *M* (*t, x, v*) and *N* (*t, x, v*) are structured according to time *t* ≥ 0, position *x* ∈ *Ω*, where *Ω* ⊂ ℝ^*n*^, and orientation *v* ∈ *S*^*n−*1^, being *S*^*n−*1^ the unit sphere. *B*(*t, x*) and *R*(*t, x*) vary only with time and position. Note that within the present iteration of the model we exclude the possibility of glioma cell invasion outside the TMT network, i.e. we do not allow movement of unconnected glioma cells. This restriction allows us to focus on the extent to which dissemination of cells via a developing TMT network can drive expansion of the tumour.

**Fig. 1.**
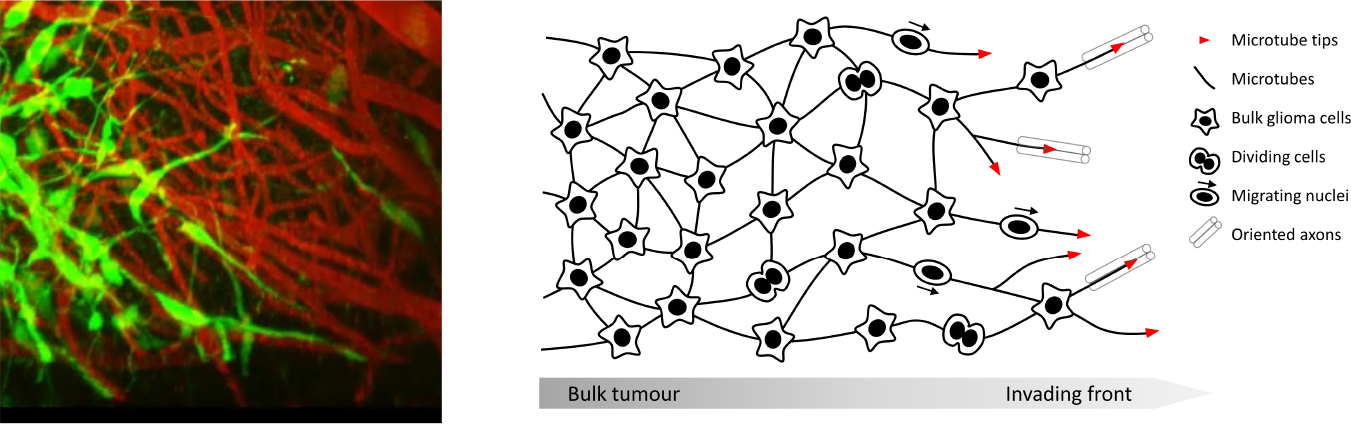
Left: Microscopic image of invading glioma (green) in to healthy brain tissue (red). The microtubes are clearly visible at the leading edge and as connections between glioma bulk cells (taken at day 23 from Osswald’s video [28], with permission). Right: Schematic showing features of microtube network growth and model variables. Microtubes sprout from bulk glioma cells, invading healthy tissue and infiltrating along aligned structures. Bulk tumour cells divide, with new nuclei subsequently capable of migrating along microtubes before settling and transitioning to mature glioma cells.

**Fig. 2.**
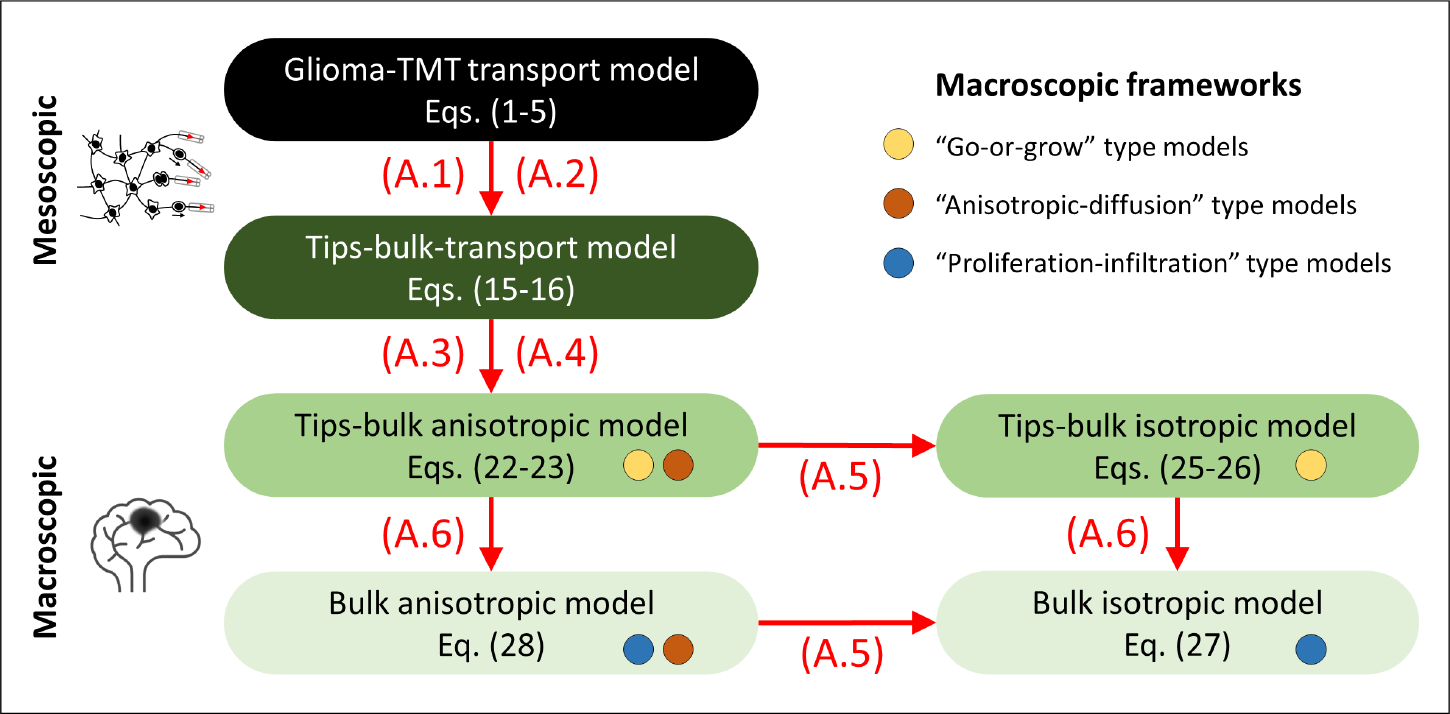
Models formulated in this work and their relationship to other models in the literature. We formulate a detailed model to describe the TMT-network driven growth of a glioma, at the mesoscopic scale. Simplifying assumptions (A.1,A.2) reduce this to a two variable model for tips and bulk, also at a mesoscopic scale. Scaling (A.3,A.4) allows translation to a macroscopic scale, resulting in an anisotropic-diffusion model for tips and bulk of go-or-grow nature. This model can be reduced further through (A.5) to an isotropic go-or-grow model (two variables) or through (A.6) to an anisotropic proliferation-invasion model (one variable). From each of these, subsequently applying the alternative assumption reduces to the simplest model in our hierarchy, an isotropic PI model.

The model equations are given by

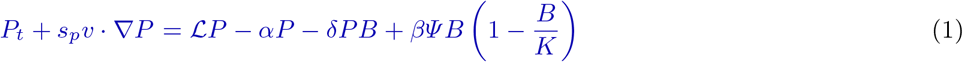

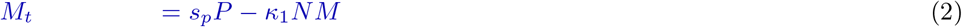

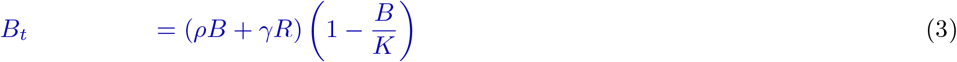

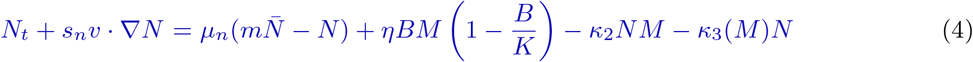

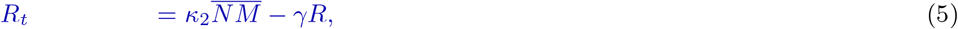

We now discuss in detail the (modelling) assumptions for each variable, referring to the observations (O1-7) listed above. A summary of variables and parameters is given in Table 1. We also list there a set of base-level values, the rationale for which will be reserved for Section 4.1.

**Table 1.** Model variables and parameter values, their meaning and units, and a selection of base values.

| Variable | meaning | initial values | units |
| --- | --- | --- | --- |
| $P(t, x, v)$ | TMT tip density (orientation $v$) | 0 | #tips/mm<sup>3</sup> |
| $M(t, x, v)$ | TMT network density (orientation $v$) | 0 | #mms of TMTs/mm<sup>3</sup> |
| $B(t, x)$ | bulk density | 0.1 K | #cells/mm<sup>3</sup> |
| $N(t, x, v)$ | migrating nuclei density (orientation $v$) | 0 | #nuclei/mm<sup>3</sup> |
| $R(t, x)$ | pre-bulk density | 0 | # cells/mm<sup>3</sup> |
| ODE Parameters | meaning | base values | units |
| $\alpha$ | tip loss rate | 0.2 | 1/hr |
| $\beta$ | rate of tips produced per bulk cell | 0.6 | *tip units/(bulk units · hr)* |
| $\delta$ | TMT to bulk connection rate | 10<sup>-5</sup> | 1/(bulk units · hr) |
| $\kappa_1$ | TMT conversion to pre-bulk | 2 × 10<sup>-6</sup> | 1/(nuclei units · hr) |
| $\kappa_2$ | nuclei conversion to pre-bulk | 2 × 10<sup>-6</sup> | 1/(network units · hr) |
| $\kappa_3(M)$ | nuclei stopping in absence of MT | 0.1 exp(-$M(t, x, v)$) | 1/(hr) |
| $\rho$ | bulk to bulk division (direct growth) | 6 × 10<sup>-4</sup> | 1/hr |
| $\eta$ | bulk to nuclei division (indirect growth) | 10<sup>-6</sup> | 1/(network units · hr) |
| $K$ | bulk carrying capacity | 25000 | cell units |
| $\gamma$ | pre-bulk maturation rate | 0.02 | 1/hr |
| $\varphi$ | Ratio of tips over bulk cells | 1.6 | 1 |
| Spatial Parameters |  |  |  |
| $s_p$ | tip invasion speed | 0.0048 | mm/hr |
| $s_n$ | nuclei movement speed | 0.0028 | mm/hr |
| $\mu_p$ | tip turning rate | 1.8 | 1/hr |
| $\mu_n$ | nuclei turning rate | 0.01 | 1/hr |
| $\nu$ | convex combination parameter | 1/2 | 1 |

1. **TMT tips** *P* (*t, x, v*), are measured as a density (i.e. number per unit volume of physical and velocity space). The tips move through space with constant speed *s*_*p*_ *>* 0 in a somewhat random “probing” fashion (O2), but with a directional bias according to the anisotropic tissue structure (O5). The oriented structure of the background tissue is encoded within a distribution *q*(*x, v*), which for simplicity is assumed to be time-invariant. Tips are further assumed to have a tendency to grow in a straightline direction, which we model by a persistent random walk as governed by a velocity-jump process [31]; we denote the corresponding equilibrium distribution as *Ψ* (*v*). Equation (2) is stated in terms of the kinetic-transport equation description, see Appendix A, where we also define *Ψ* (*v*) explicitly in (34). Noting that TMTs frequently retract (O2), but can also emerge from existing TMTs (O3), we consider an effective loss rate *α* as the difference: *α* can therefore be positive or negative. Tips are generated through sprouting from bulk cells, but can also be lost through connecting with bulk cells (anastomosis, O6). Anastamosis is assumed to arise on contact between tips and bulk cells, with a rate *δ* per bulk cell. The logistic form for forming new sprouts from bulk cells provides a saturation on the formation of new network as the tumour reaches its carrying capacity. The growth rate of new sprouts is *β ≥* 0, oriented in directions according to the tissue structure. As we aim at recovering an aggregate, or *macroscopic*, description of the system, we introduce the macroscopic density of the tips as

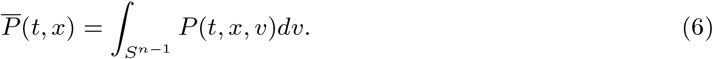
2. **TMTs** *M* (*t, x, v*), are measured as the number of units of lengths of TMTs per unit volume with orientation *v*, where we have adopted the “snail-trail” framework [10] to assume that TMT tip movement leads to the formation of microtube in its wake. Loss of the microtube occurs as nuclei migrating along the network (O4) come to a rest and transition, with a section of microtube, into pre-bulk cells. For simplicity we assume this occurs at a rate proportional to *NM* . Analogously to as stated above for the tips, we define the macroscopic density of the microtubes

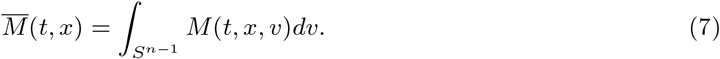
3. **Bulk** *B*(*t, x*), is assumed to grow until reaching a carrying capacity *K >* 0. Bulk growth occurs either through direct bulk cell division, at a rate *ρ >* 0, or through the maturation of pre-bulk cells into bulk, at a rate *γ >* 0. Note that we assume there is no movement/transport of bulk tumour cells, except through the below described dissemination of nuclei along the microtube network. We also assume that the TMT tips, TMTs, and nuclei carry essentially no volume, i.e. they do not contribute to the carrying capacity of the bulk.
4. **Migrating Nuclei** *N* (*t, x, v*), are generated in some proportion of bulk divisions, with at least one of the divided nuclei travelling along the TMT network and settling at a point distant from the division site. As for tip movement, their dynamics are described in terms of a kinetic-transport equation. Nuclei are assumed to move with a characteristic movement speed *s*_*n*_, and with a direction of movement chosen from the (normalised) distribution of the microtube network, i.e.

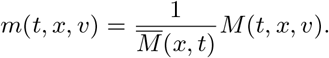

Nuclei stop moving based on two processes. There is a spontaneous stopping with rate *κ*_2_ along tumour microtubes. They will also stop moving once they reach the end of a microtube. We describe this here with a microtube-density dependent removal rate *κ*_3_(*M*), which is assumed to decrease to negligible levels when the microtube network density is high. The macroscopic density of the nuclei is defined as

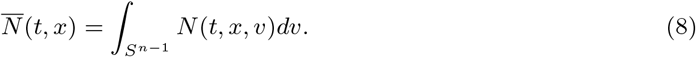
5. **Pre-bulk** *R*(*t, x*), forms as migrating nuclei come to a rest and, with surrounding microtube, transition into a bulk cell. We call this a pre-bulk stage and assume that the transition to pre-bulk occurs with rate parameter *κ*_2_ and according to the local microtube density. Since the pre-bulk has no movement orientation, we use the integrated terms

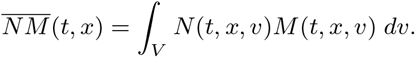

The pre-bulk matures into bulk, with rate *γ*, at which point it can start undergoing mitosis. We note, though, that if the bulk is at capacity then the pre-bulk simply decays.

Unspecified in (1) is the choice of the turning operator ℒ, that describes the reorientation of TMT tips as they extend through brain tissue. This operator is taken to be of general form

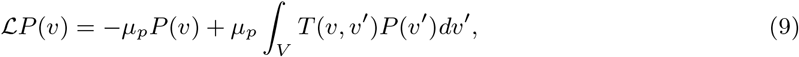

where *µ*_*p*_ is the rate that the tips turn and *T* (*v, v*^*i*^) defines the turning kernel. Here we ignore the dependence on space *x* and focus on the *v* dependence of the turning terms. In our full model the terms below *q, T*_1_, *T*_2_, *T* will all depend on space *x*.

Noting that TMTs have both a tendency to follow the direction of nerve fibres and maintain a relatively straight line (i.e not bending excessively), we choose a turning kernel that combines a path-following term, denoted by *T*_1_, and a persistent random walk, denoted by *T*_2_. Specifically,

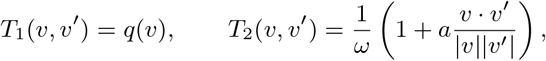

where *ω* = |*V* | and 0 *< a <* 1 is a persistence parameter ([17, 32]. The function *q*(*v*) describes the orientation response to aligned structures (e.g. nerve fibres, capillaries etc) and can potentially be informed through DTI imaging [42]. As described more fully in Appendix A, the turning kernel is taken to be a convex combination of *T*_1_ and *T*_2_:

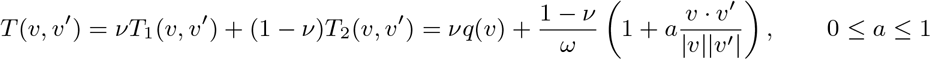

where the parameter *ν ∈* [0, 1] is the anisotropy parameter, which reflects the weighting between persistence and aligned structures on the orientation.

In terms of nuclei movement, the choices in (4) stipulate that nuclei move with speed *s*_*n*_, change direction with rate *µ*_*n*_, and assume the reorientation at a turn follows the angular distribution of the mature TMT network.

Finally, we note that (1)-(5) must be equipped with suitable boundary conditions. The brain sits inside the skull, which is (mostly) rigid. However, with the analysis presented here at the micrometer to milimeter scale we can conveniently assume that the tumour origin is sufficiently far away from the skull. This in turn allows us to assume an (effectively) infinite domain and avoids the need to specify boundary conditions. If the tumour originates closer to the skull or another boundary, other conditions may be necessary.

### 2.3 Analysis of the Kinetic Part

To initiate a deeper understanding of the dynamics of model (1-5), we begin with an analysis of the kinetic terms, i.e. the time-only dynamics for a system that excludes spatial terms. In other words, we assume all variables *P, M, B, N* and *R* are constant in space and velocity, and only depend on time. Certain terms in equations (1-5) contain integrated variables, for example

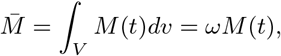

which generates an additional parameter *ω* = |*S*^1^| = 2*π* in 2-D and *ω* = |*S*^2^| = 4*π* in 3-D. Integrating and introducing 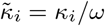 for *i* = 1, 2, 3. We have

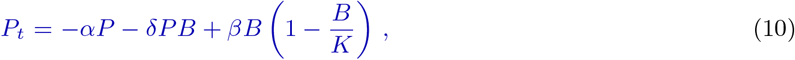

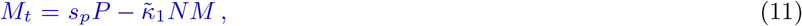

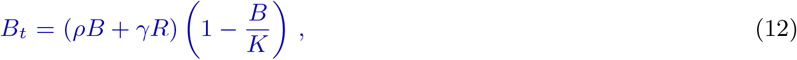

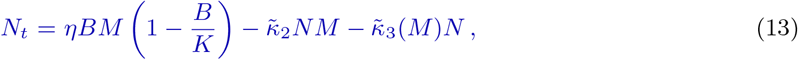

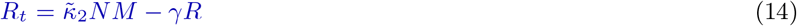

where we have dropped the 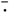 over *P, M* and *N* as the only dependence in both cases is on *t*. The only steady states of the above system are part of two continua of steady states: the set of disease-free equilibria

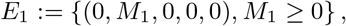

and the set of diseased equilibria

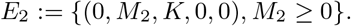

We note that in both cases the density of tumour microtubes is a free parameter. The value *M*_1_ indicates that a certain microtube residue can persist and will not be removed. The parameter *M*_2_ is an effective carrying capacity for the TMT, which depends on the initial conditions and the time dynamics of the entire system.

Linearizing the above model about the steady states, we find the following Jacobian for the disease-free state

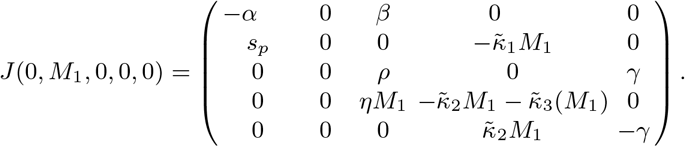

When *M*_1_ is zero we obtain eigenvalues *−α, −γ*, 0, 0, *ρ*, indicating that the disease free state is unstable. For general *M*_1_ *>* 0 we find two eigenvalues (*λ*_1_ = *−α, λ*_2_ = 0) and three further by solving the cubic equation

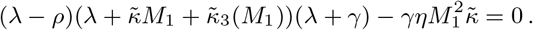

Noting that this cubic has positive leading order term and negative vertical intercept, it must have at least one positive real root. Consequently, the disease-free state is unstable. In particular, it appears that introducing a small bulk with or without a small residual of tumour microtubes is sufficient to induce movement away from the disease-free state and initiate tumour growth.

The Jacobian for the diseased state is

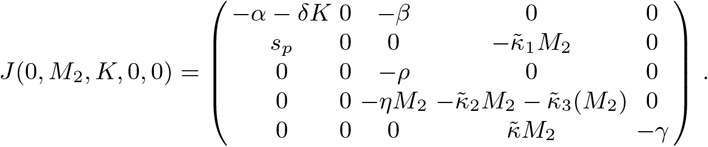

Here all five eigenvalues are directly evaluated as 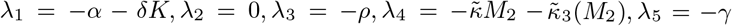 and the diseased state is stable, except for the eigenvalue 0 that corresponds to the continuum of steady states parameterised by *M*_2_. Therefore, we expect growth of the tumour that eventually settles on some diseased state within *E*_2_.

### 2.4 Dynamics of the Glioma-TMT Transport Model

To demonstrate that the model captures essential features of glioma-TMT dynamics, we perform some preliminary simulations. Note that a more detailed suite of numerical simulations will be performed later (in Section 4), where we also provide details of the model parameters; where possible, these are derived from the observations of Osswald et al. [29], see Table 1.

First we consider the temporal dynamics by simulating the ODE model (10-14), see Figure 3. We observe in Figure 3 A) that early dynamics reveal a quick growth of TMT tips (red), followed by a steady growth of the TMT network (blue). The growth of bulk (black), migrating nuclei (tan) and pre-bulk (purple) follows after some delay. As the bulk approaches the carrying capacity (Figure B)), the mature TMT network converges towards a steady state (*M*_2_), with the components (*P, N, R*) converging to zero.

**Fig. 3.**
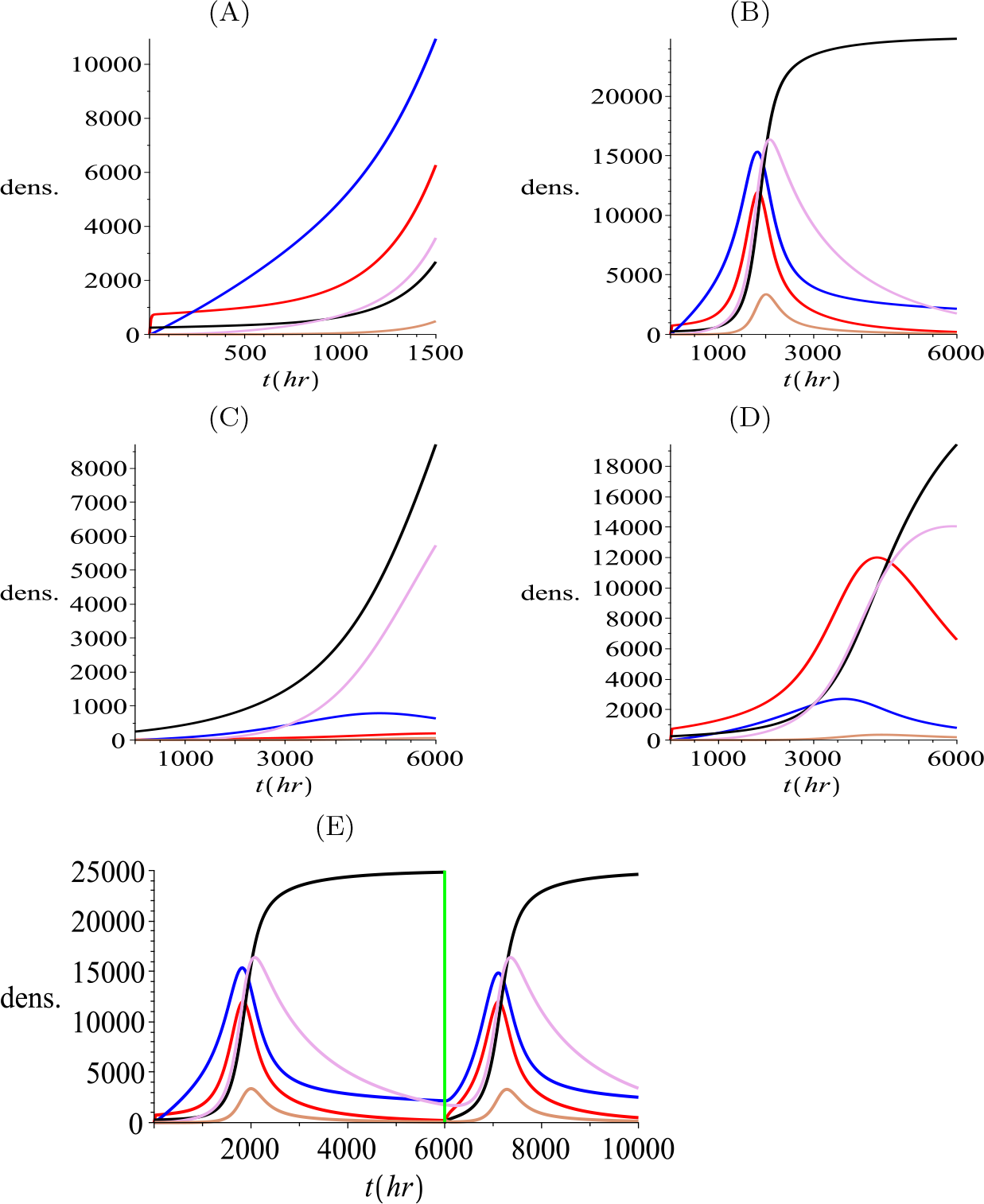
Simulation of the ODE model (10-14), showing tips (*P* red), TMTs (*M* blue), bulk cells (*B* black), nuclei (*N* magenta), and pre-bulk (*R* brown). Unless stated otherwise, parameter values are chosen according to the reference set listed in Table 1. A) Initial evolution up to time *t* = 1500 hours. B) Dynamics up to time *t* = 6000 hours, showing approach to the steady state. C) Dynamics under reduced tip production (*β* reduced to 0.01. D) Reduced MT growth rate (*s*_*p*_ reduced to *s*_*p*_ = 0.00048). E) Simulated treatment in which bulk cells are removed at time *t* = 6000 hours (vertical green line), but microtube network is left intact. For all simulations, initially we set *P* (0) = 0, *M* (0) = 0, *B*(0) = 0.01*K, N* (0) = 0, *R*(0) = 0.

Variations of parameters about the default parameters indicate that the system is most sensitive to the TMT growth rate *s*_*p*_ and the tip production rate *β*. Osswald et al. [29] indicate that the TMT dynamics could provide new treatment targets for chemotherapy. We test this here in the ODE model by reducing the tips production rate by a factor 60 (in Figure 3 C)) and the TMT growth rate *s*_*p*_ by a factor 10 (in Figure 3 D)). Reducing the sizes of these parameters leads to significantly delayed tumour growth and a mature TMT network density that settles to a lower value at equilibrium.

Notably, these results are consistent with observations in [29], where engineered GBMSCs (glioblastoma multiforme stem cells) with a genetic knockdown of GAP-43 (a protein relevant for the formation of neurite-like membrane protrusions) results in a lower speed of invasion, structurally abnormal TMTs with reduced branching, smaller proliferation capacity and, eventually, a marked reduction of the tumour size. In terms of other parameters, we found that varying the pre-bulk to bulk maturation rate *γ* did not substantially change the overall dynamics, but an increase in *γ* would reduce the size of the pre-bulk population (simulations not shown). Increasing the bulk growth rate *ρ*, unsurprisingly, leads to faster overall growth and a reduction in the delay between TMT network growth and bulk growth. Increasing the rate of nuclei division *η* leads to an unrealistically large moving nuclei component, while variation in the rate of nuclei transport (*s*_*n*_) along TMTs had relatively little impact on the dynamics. A common feature of all simulations is that tips and TMTs lead the growth, followed by the bulk and the nuclei. Eventually *P, N, R* converge to zero and *M* and *B* settle at a non-zero equilibrium, consistent with the predictions of the steady state analysis.

In Figure 3 E) we simulate a treatment at time *t* = 6000, where all bulk tumour cells are removed but all other components are left behind. We observe that a new tumour soon develops following this procedure, which grows from nuclei and pre-bulk integrated within the remaining TMT network. This implies that it would be important to also remove any invading TMT network for successful treatment. We proceed to explore spatiotemporal aspects of growth, concentrating on a one-dimensional setting that represents a transect line through the tumour, see Figure 4. Here, an initial tumour mass is seeded at the left of the domain, with the right portion therefore representing non-diseased brain tissue. Simulations suggest the emergence of travelling wave/pulse profiles, i.e. where model variables evolve to profiles with constant shape and moving into the non-diseased tissue with constant speed. The wavefronts for tips (red), microtubes and nuclei (magenta) are a little ahead of the bulk (black) and pre-bulk (tan) waves. This clearly shows the invasive nature of microtubes, which provide channels for the migrating nuclei to infiltrate and seed new bulk. In the rear of the wave, the tumour bulk reaches a carrying capacity while the free tips and the migrating nuclei approach zero. The microtubes settle on a non-zero steady state value, consistent with observations of an established network of microtubes that interlink glioma cells in the tumour mass.

**Fig. 4.**
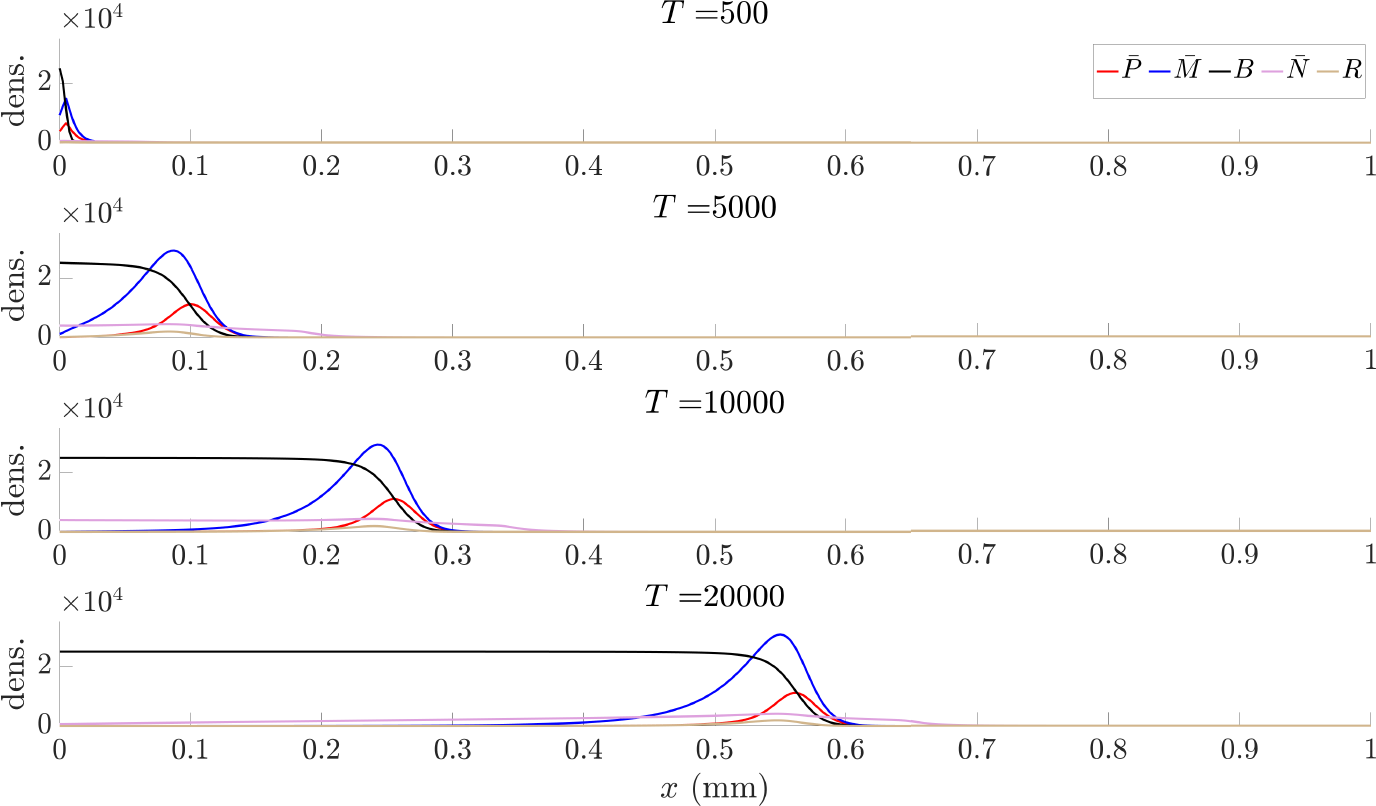
Illustrative dynamics of the full glioma-TMT transport model (1-5) for a quasi-1D setting, describing invasion into a portion of tissue of length 1 mm and characterised by an isotropic fibre arrangement. Initially we assume the bulk is described by a half-Gaussian, with maximum density at the origin and equal to the carrying capacity (*B*(*t* = 0, *x* = 0) = *K* = 30) and standard deviation 0.05 mm. All other variables are initialised as zero. Each frame plots the distributions at (top to bottom) 500, 1000, 10000 and 20000 hours. Other initial condition with tumour seed on the left of the domain show a very similar invasion process (simulations not shown). Parameter values as in Table 1.

## 3 Model Reductions

As mentioned earlier, the spatial spread of glioma has formed the focus of many modelling studies [43, 44, 33, 11, 42, 40, 7], with many of these falling into a classical reaction-diffusion framework. With the aim of reaching a model that is of similar tractability, we perform a sequence of model reductions. The first step is a quasi-steady state assumption for the (non tip) TMTs and nuclei, which directly leads to models of “go-or-grow” type, popular in various glioma modelling studies [12, 3, 34, 15, 13, 38, 40]. Further scaling leads to “fully anisotropic” models, which have been explored in a number of recent studies [33, 11, 42] and aim to account for the impact of tissue anisotropy on glioma growth. Finally, an assumption of isotropy leads to the prominent proliferation-infiltration type models, e.g. [43, 44, 36, 22], which fall into the classical class of Fisher-KPP equations, and therefore admit straightforward deduction of the wave speed. Within each simplifying step we stress the underlying assumptions, clarifying the biological processes retained and those that are neglected. Later, a numerical analysis is used to investigate (quantitatively) the loss of information as we proceed through the reductions. A schematic showing the step-by-step sequence of assumptions as we proceed through the reductions is provided in Fig. 2, and a summary of these scaling arguments is provided in Section 3.5.

### 3.1 Reduction to Go-or-Grow Type Models

As the first step, we consider the following set of assumptions. **Assumptions (A):**

**(A.1)** We assume that the dynamics involving nuclei and pre-bulk are in quasi-equilibrium.

**(A.2)** Microtubes are assumed to be in quasi-equilibrium and follow the directions of the tips.

In other words, we consider equations (2), (4) and (5) to be in quasi-equilibrium. Then (5) in equilibrium implies

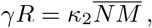

and from (2) we have

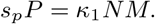

Taken together,

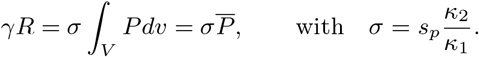

Using this relation in the bulk equation (3), the system for (*P, B*) is observed to decouple from the remainder and we obtain the *tips-bulk-transport* model:

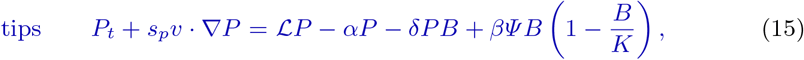

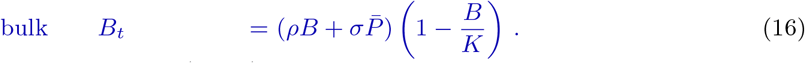

An inspection of the terms in (15-16) reveals that bulk grows but it does not move spatially, while the tips explore the environment via a correlated random walk. This tip expansion leads (via the assumed instantaneous processes of microtube extension, nuclei movement, transition to pre-bulk and maturation to bulk) to the effective seeding of new bulk and overall tumour spread. Model (15-16) is therefore essentially of a *go-or-grow* structure [12, 3, 34, 15, 13, 38, 40], with the difference being that typical go-or-grow models are compartmentalised into two cell sub-populations (an actively-invading compartment and an actively-proliferating compartment) while the invaders here are not cells but TMT tips. Nevertheless, the go-or-grow type model arises naturally from the microtube invasion model. Moreover, with respect to the fact that here the invaders are TMT tips, we remark that in our model invading cells move by means of the TMT network and that, thanks to assumption (A.2) the invading tips are proportional to the product of the TMT density and of the nuclei of the invading cells.

### 3.2 Parabolic Scaling

A parabolic scaling is used to separate time scales. Specifically, rather than considering a microscopic timescale of seconds and a spatial scale of micrometers, we transform to macroscopic scales of hours and centimeters. For the parabolic scaling we make the following assumptions.

**(A.3)** A rescaling of time and space to the macroscopic scales *τ* = *ε*^2^*t, ξ* = *εx*, where *ε <<* 1 is a small parameter.

**(A.4)** Reaction terms scale as *ε*^2^, i.e. reaction terms are slow.

The two assumptions (A.3) and (A.4), applied to (15), lead to

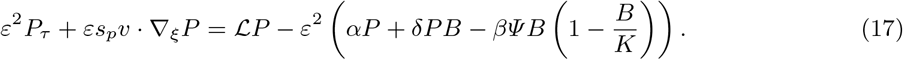

We assume the tip density *P* (*τ, ξ, v*) can be written as a regular expansion in *ε*

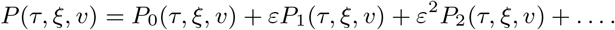

and we substitute this expansion into the scaled equation (17), collecting terms of equal order in *ε*. The leading order terms *ε*^0^ are

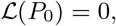

Which implies that *P*_0_ *∈* ker ℒ. We computed the null space of ℒ in Appendix A in Lemma 1, thus

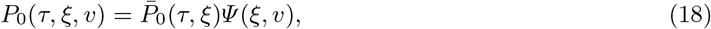

where 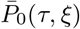 depends on space and time but not on velocity. The terms of order *ε*^1^ are

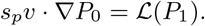

At this point, we posit that *P*_1_ *∈ (Ψ)*^*⊥*^. On this space we can invert ℒ (see [17]), yielding

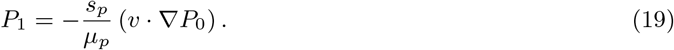

Indeed, *P*_1_ *∈ (Ψ)*^*⊥*^, since

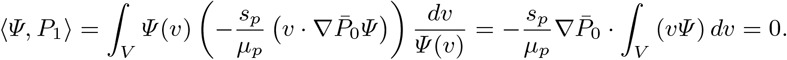

Where the last integral is zero since *Ψ* (*v*) from (34) is symmetric in *v*. Next, we look at the terms of order *ε*^2^:

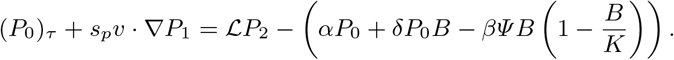

Integrating this equation over *V* and substituting *P*_0_ from (18) and *P*_1_ from (19) yields a macroscopic system that can be combined with the bulk equation (16):

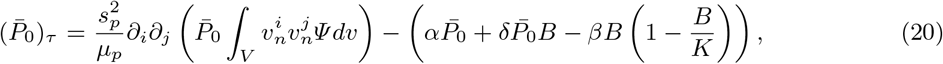

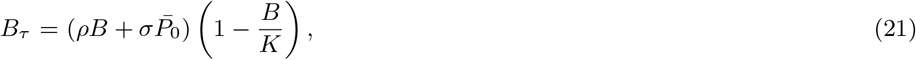

where we use summation convention on repeated indices. Note that several details within this calculations have been omitted, since they are standard and can be found elsewhere (for example, see [17]). We can simplify the notation by setting the leading order term as 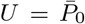 and using tensor notation for the diffusion term. Specifically,

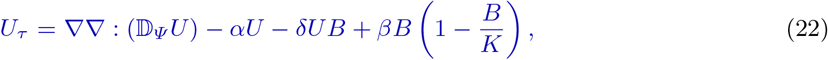

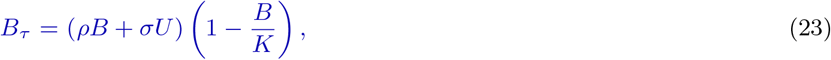

with diffusion tensor

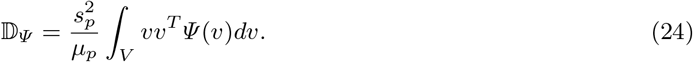

We remark that the system (22-23) is again a model of go-or-grow type, but here the movement term is an anisotropic diffusion term. We refer to (22-23) as the *tips-bulk-anisotropic* model.

### 3.3 The Isotropic Case

In [32] we classified choices for *Ψ* (*v*) such that the diffusion tensor in (24) is isotropic, i.e. proportional to the identity matrix. One such case occurs when *Ψ* depends only on the speed *s*_*p*_, not on the direction: *Ψ* (*s*_*p*_*v*) = *Ψ* (*s*_*p*_). Then, Δ_*Ψ*_ = *d*𝕀 for some constant *d >* 0. This generates the next assumption.

**(A.5)** We assume Δ_*Ψ*_ = *d*𝕀 for some constant *d >* 0.

Consequently, we obtain the *tips-bulk-isotropic* model

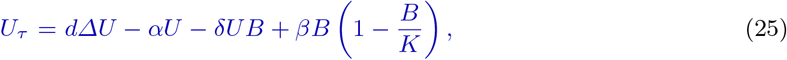

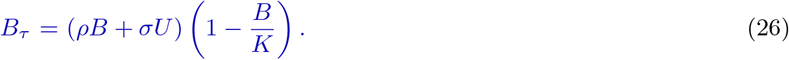

This model (25, 26) is the standard go-or-grow model, which has been studied in several publications [3, 34, 15, 40].

### 3.4 A Single Equation for the Bulk

To obtain an equation for the bulk alone, we make the natural assumption that, on average, the number of tips per bulk is constant, i.e.

**(A.6)** *U ≈ ϕB*.

We use parameter estimation in Section 4.1 to estimate *ϕ*. Now consider the time derivative of the sum of equations (25, 26) and obtain

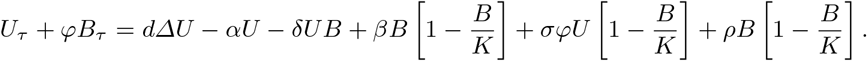

Using the above assumption (A.6), we can write this in terms of *B* alone obtaining the *isotropic bulk model*,

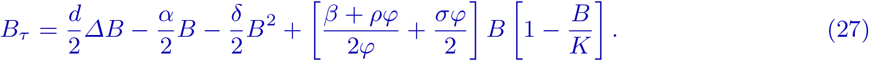

The same approximation can be performed for the anisotropic model (22-23) and we get

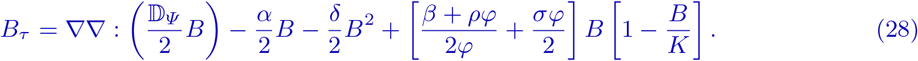

We note that the above model has been used in, for example [42, 33, 11], to model glioma growth based on diffusion-tensor imaging data.

The equation (27) is simply a standard Fisher-KPP model with isotropic diffusion and a monostable forcing function. In particular, we can write (27) as

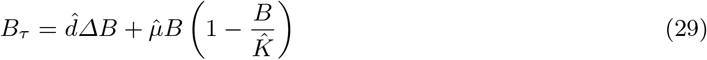

with

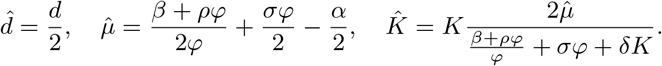

Within the glioma modelling literature, model (29) is referred to as the PI-model (e.g. [44, 22]). Note that earlier iterations of the PI-model used an exponential growth term for the proliferation, while others have favoured logistic growth (e.g. [44, 42]). Here we have shown how the PI model can be derived from models for microtube-network driven expansion. The logistic growth in the macroscopic model (29) is obtained by assuming a logistic growth of both the tips and the bulk at the mesoscopic level in (1) and (3).

The formulation as a Fisher-KPP model (29), of course, allows direct computation of a minimal invasion speed:

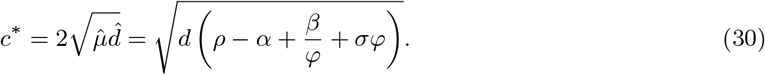

We note that the invasion speed (30) increases with the tip production rate *β*, the bulk growth rate *ρ*, and in the diffusion coefficient *d*. It decreases for increasing tip removal rate *α*.

### 3.5 Summary of Model Reductions

To conclude this section we summarize the model reductions and the corresponding models. Figure 2 lists the series of models, with the successive assumptions

**(A.1)** Nuclei dynamics are in quasi-equilibrium

**(A.2)** Microtubes are in quasi-equilibrium with the tips.

**(A.3)** Rescaling of macroscopic time and space scales *τ* = *ε*^2^*t, ξ* = *εx*, where *ε <<* 1.

**(A.4)** Reaction terms scale like *ε*^2^.

**(A.5)** Isotropy Δ_*Ψ*_ = *d*𝕀 for some constant *d >* 0.

**(A.6)** Tips and bulk are proportional; *U ≈ ϕB*

## 4 Numerical Comparisons

### 4.1 Parametrization

As units, we use a spatial scale of millimetres (*mm*) and measure time in hours (*h*). Our *baseline* parameter set, reported in Table 1, is estimated according to the various data and videos reported in [28, 29]. Consequently, the parameters described here are linked to the associated experimental set-up (transplantation of patient-derived primary brain tumour cells to the mouse brain) and would require reevaluation in the context of, say, predicting glioma evolution in a patient.

Concerning the kinetic parameters that drive TMT dynamics, video data in [29] indicate that new tips emerge from a cell approximately once every 100 minutes, leading to *β* = 0.6 */hr*. Anastomosis is assumed to be relatively rare where, if we assume that only one in 20 tips undergoes anastomosis, we take *δ* = 0.05*β* = 0.03 */hr*. Further observations of videos [28] indicate that approximately only 20% of tips lead to mature microtubes, hence *α* = 0.2 */hr*.

For the parameters that drive cellular (bulk and nuclei) kinetics, the videos from Osswald [28, 29] suggest that the bulk doubles every *∼*20 days, i.e. a net bulk growth rate of about 1 *×* 10^*−*3^ */hr*. In our model, bulk growth stems from two effects: the maturation of nuclei that have moved along a TMT, and through direct mitosis of bulk cells. Noting that direct mitosis is rarely observed in these experiments, we chose *ρ* = 6 *×* 10^*−*4^*/hr*, which corresponds to a doubling time for bulk alone (without TMT) of about 150 days.

The transition of pre-bulk into bulk cells is taken to be approximately 5 hours, i.e. *γ* = 0.2 */hr*. Parameters *η* and *κ*_1_, *κ*_2_ dictate the generation of new migrating nuclei and their subsequent conversion into resting nuclei: for the former we assume *η* = 8 *×* 10^*−*4^ */*(*cell units · hr*), and for the latter we take *κ*_1_ = *κ*_2_ = 0.1 */*(*cell units · hr*). The term that removes migrating nuclei in the absence of microtubes is given the form of a decaying exponential, *κ*_3_(*M*) = 0.1 exp(*−M* (*t*)); this stipulates the half life of nuclei of 7 hours when *M* = 0 and that negligible loss occurs for a substantial (i.e. *M »* 0) microtube network. Experimental findings are reported from image sections with a length scale *∼* 0.1 *mm*. Counting mature bulk cells in these images ([29] Fig. 1e, Fig. 4c, Fig. 4d, and day 23 in video [28]) leads to a carrying capacity of *K* = 25000 *cells /mm*^3^. We note that under the above described parameters, the simulations reported earlier under kinetic-only terms indicated a maximum doubling time for bulk cells in the region of 10-20 days (see Figure 3).

Regarding parameters that govern movement dynamics, time lapse videos in [29] of a single invading cell over 104 minutes indicate a tip (substantially) changing direction about 3 times. This leads to a tip turning rate of *µ*_*p*_ = 1.8 */hr*. Similarly, nucleus movement indicated a single change of direction in 3 days. Hence we chose the turning rate of moving nuclei to be *µ*_*n*_ = 0.01 */hr*. The videos also allow calculations for the invasion speeds, which we estimate at *s*_*p*_ = 0.0048 *mm/hr* for the TMT tips. The nuclei invasion speed is reported in [29] as *s*_*n*_ = 0.0028 *mm/hr*.

We determine the proportionality constant *ϕ* in assumption (A.6) from data in [29]. This constant describes the average number of microtubes per bulk cell; Figure 2 in [29] provides a histogram for the number of protrusions per cell as the tumour progresses, and we take the mean at day 20 to obtain *ϕ* = 1.6.

The parameters estimated here are summarized in Table 1. Note that in Figure 3 and Figure 4 we provided simulations for the model under these parameters, respectively for the kinetics-only system and of the full system for a one dimensional invasion wave scenario.

### 4.2 Numerical Test Cases

Here we present the results of numerical simulations in two dimensions for the full microtube model (1-5), along with the anisotropic diffusion approximations (25-26) and (28). Regarding the numerical methods, the equations that involve transport terms in (1-5) are integrated with the methods developed in [26, 25]. The remaining equations are either of reaction-diffusion type or ODEs, and they are integrated with an Explicit Euler time integration scheme. For the anisotropic and isotropic diffusion models obtained following the parabolic scaling, we use an adapted method from [45]. Simulations are performed on bounded rectangular regions, where boundary conditions are imposed in a manner that leads to zero flux at the macroscopic level.

Numerical simulations are performed under three scenarios, designed to explore how quantitative and qualitative solution properties change under the various model formulations. Unless stated otherwise, parameters are taken from the baseline set presented in Table 1. Configurations are as follows:

**Test 1 (invasion wave)**, exploring a one-dimensional invasion process within an isotropic tissue;

**Test 2 (anisotropic stripe)**, exploring a two-dimensional invasion process for a tumour initialised within a stripe of highly aligned fibres;

**Test 3 (treatment)**, exploring a simplified treatment scenario in which all but a small fraction of bulk cells are removed and the possibility of cancer recurrence is investigated.

#### 4.2.1 Test 1: Invasion Waves

We consider a quasi-1D setting in which the domain is elongated along the *x*-axis and solutions are constant in *y* direction. A tumour is initialised as bulk at the edge of a portion of tissue of length 1 mm. We note that the fibres are considered to be isotropic; hence, model (22-23) is identical to (25-26), and model (28) is identical to (27). The invasion process for the full model (1-5) has previously been shown in Figure 4.

In Figure 5 we compare solutions at times *T* = 500, 1000, 20000 hours, for each of the full model (1-5), the tips-bulk-transport model (15-16), the tips-bulk-isotropic model (25-26), and the bulk-isotropic model (27). We observe that all models generate an invasion front, in which the bulk steadily infiltrates non-diseased tissue, and appears to be of travelling wave form (constant speed and profile). We notice immediately that (under the chosen parameter values) the different models yield very different invasion speeds. Table 2 lists these speeds, measured according to the location of the 20% level set for the bulk. The invasion speed increases with each scaling or simplification, resulting in more than an order of magnitude increase for the Fisher-KPP limit version when compared to the full model. Notably, the measured invasion speed from experiments (also reported in Table 2) lies between the simulated values and is of the same order of magnitude.

**Table 2.**
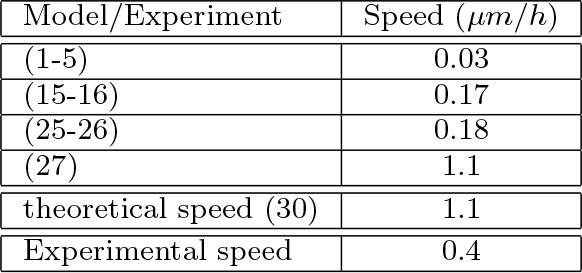
Test 1, wave speed comparison from Section 4.2.1. The table lists the propagation wave speeds of the bulk as computed for the different models, the theoretical speed from formula (30) and the measured invasion speed from video [28].

**Fig. 5.**
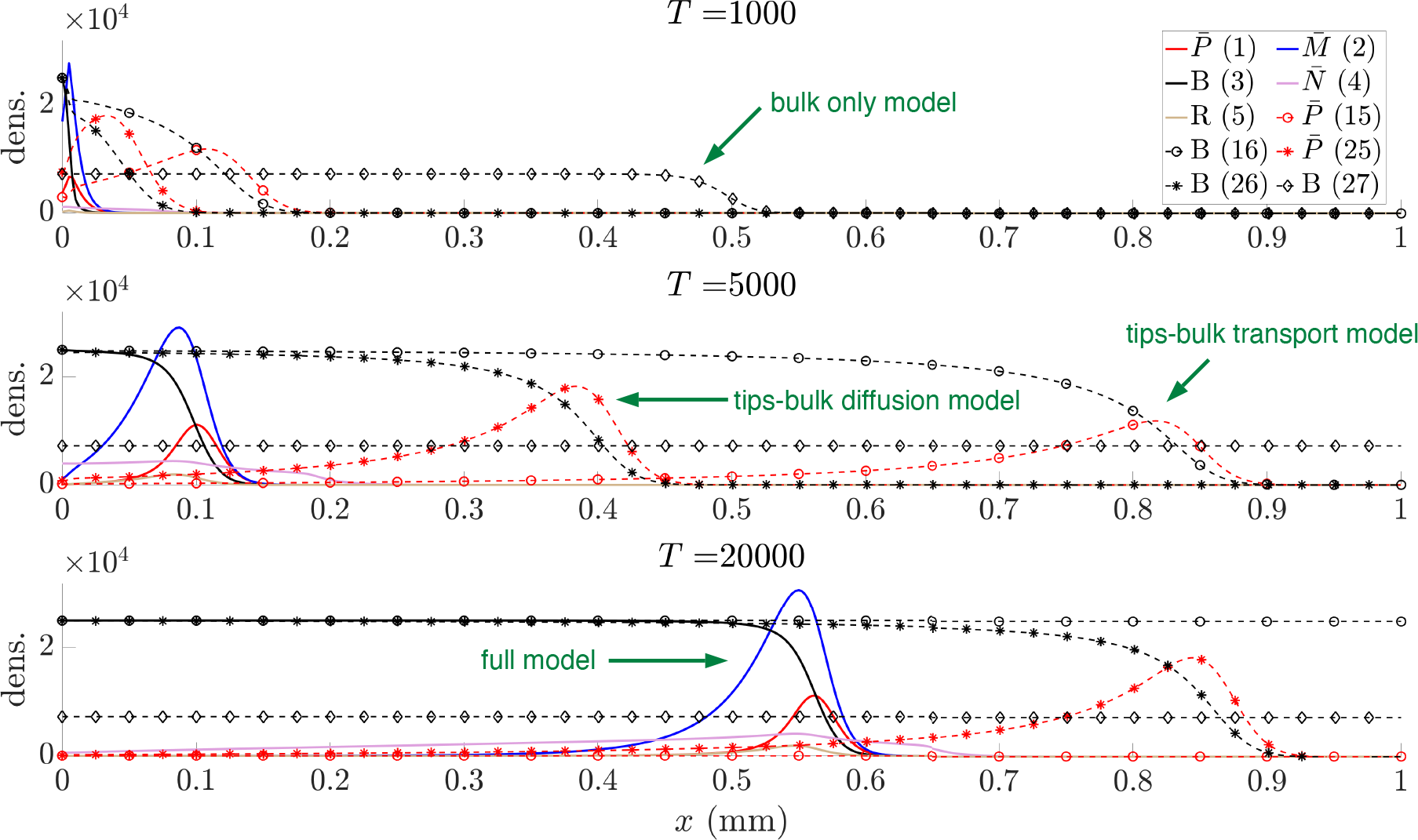
Profiles of the solutions at times *T* = 500, 1000, 20000 hours, for each of the full model (1-5), the tips-bulk-transport model (15-16), the tips-bulk-isotropic model (25-26), and the bulk-isotropic model (27). For details of the numerical schemes we refer to the text, while for parameters we refer to Table 1.

This discrepancy in the speeds of the various models is a direct result of the assumptions (A.1-A.6) made during the model reduction process, demonstrating the potential pitfall of making such assumptions without evaluation of their impact on the quantitative dynamics. In particular, the very essence of a quasi-steady state assumption is to compress the time required to arrive at a steady state. As an example, the implication of assumptions (A.1, A.2) is to instantaneously place new bulk at the position of extending microtube tips, thus ignoring the time required for nuclei to travel down microtubes and transition into bulk: the tips-bulk-transport model, therefore, overestimates the invasion speed of the full microtubes model. Moreover, derivation of the macroscopic limit utilises the diffusive scaling (A.3) and that the reaction term is assumed to be small (i.e. behaves as *E*^2^ (A.4)). Hence, the evolution as prescribed by model (26)-(25) is faster than not only (1-5) but also (15-16). On this we note [40], where the wave speeds for a go-or-grow glioma model (our model (25, 26)) and the PI-model (27) were compared analytically. It was found that the wave speeds differ by a factor that is proportional to the transition rates of the two compartments of the go-or-grow model.

Despite the disagreement at a quantitative level in the time evolution, there is a clear qualitative correspondence between the solutions: for the bulk distribution we observe that each of models predict growth of the bulk to the carrying capacity in monotonic fashion.

#### 4.2.2 Test 2: Anisotropic Scenario

We next explore a scenario that involves spatially-dependent anisotropy. Specifically, we consider a square domain *Ω* = [*−*1, 1] *×* [*−*1, 1] which contains a *stripe arrangement*, i.e. a stripe of horizontal fibres within an isotropic tissue. Reflecting the observation that microtubes tend to follow aligned structures [29], we impose a turning kernel (9) that relates tip orientation to the assumed tissue anisotropy. Following the approach of previous studies (e.g. [33, 42]), we set the function *q*(*v*) according to a bimodal von Mises distribution, parameterized by a spatially-varying direction parameter, *θ*_*q*_ *∈* [0, *π*), and concentration parameter, *k*_*q*_ *∈* [0, *∞*)

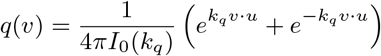

where *u* = (cos(*θ*_*q*_), sin(*θ*_*q*_))^*T*^, and *I*_0_ is the modified Bessel function of the first kind of order zero. We denote the region of horizontal fibres as *Ω*_*H*_ = [*−*1, 1] *×* [*−*0.25, 0.25] and set

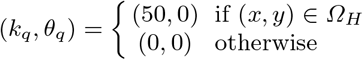

Figures 6-7 display the results of representative simulations under the stripe arrangement. In Figure 6(a) we show output for the full tumour-microtubes model (1-5). Notably, we observe an anisotropic pattern of growth, whereby the growing tumour spreads more rapidly along the stripe of aligned fibres. This is a direct consequence of the orientation and movement of tips and microtubes along the fibres, therefore biasing those directions for the seeding of new bulk. Profiles of the variables reflect those seen in the one-dimensional simulations: tips and nuclei are found to be concentrated into a pulse around the invasive front, giving rise to a combination of bulk and mature microtubes in the rear.

**Fig. 6.**
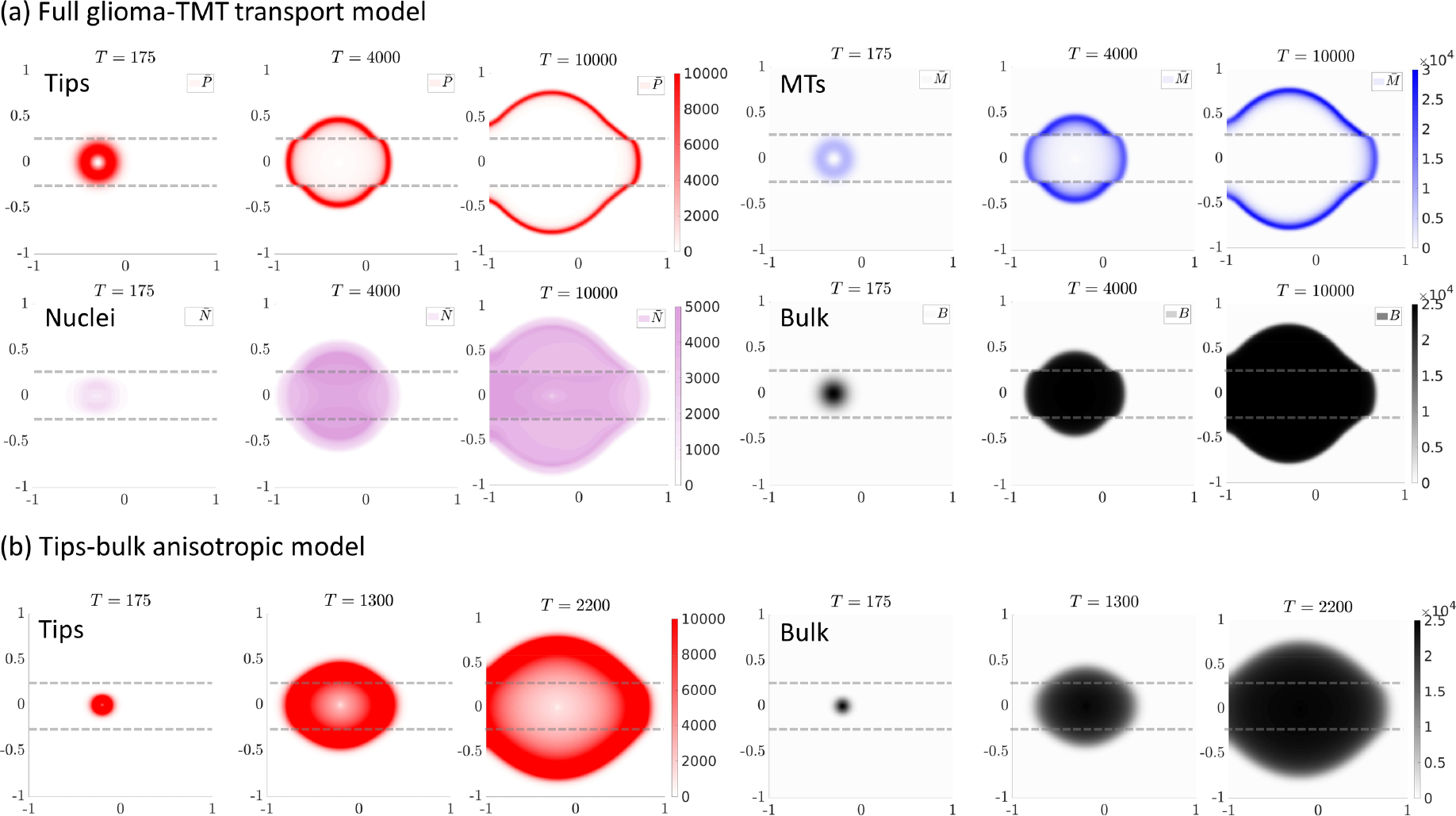
Anisotropic scenario. Representative simulations of the full tumour-microtubes model (1-5) in (a) and the tips-bulk-anisotropic model (22-23) in (b), under the stripe arrangement. The region of horizontally aligned fibres is enclosed by the dashed lines shown in each panel. For each model we show the distributions of the variables at the three times indicated above each panel. Tips are shown in red, the TMT in blue, the bulk in black and the migrating nuclei in violet. Note that for the full tumour-microtubes model, migrating nuclei and pre-bulk have similar distributions and only the former are plotted. The parameters are drawn from Table 1, where the anisotropy parameter was set to *ν* = 0.8. Length and time scales are in millimetres and hours.

**Fig. 7.**
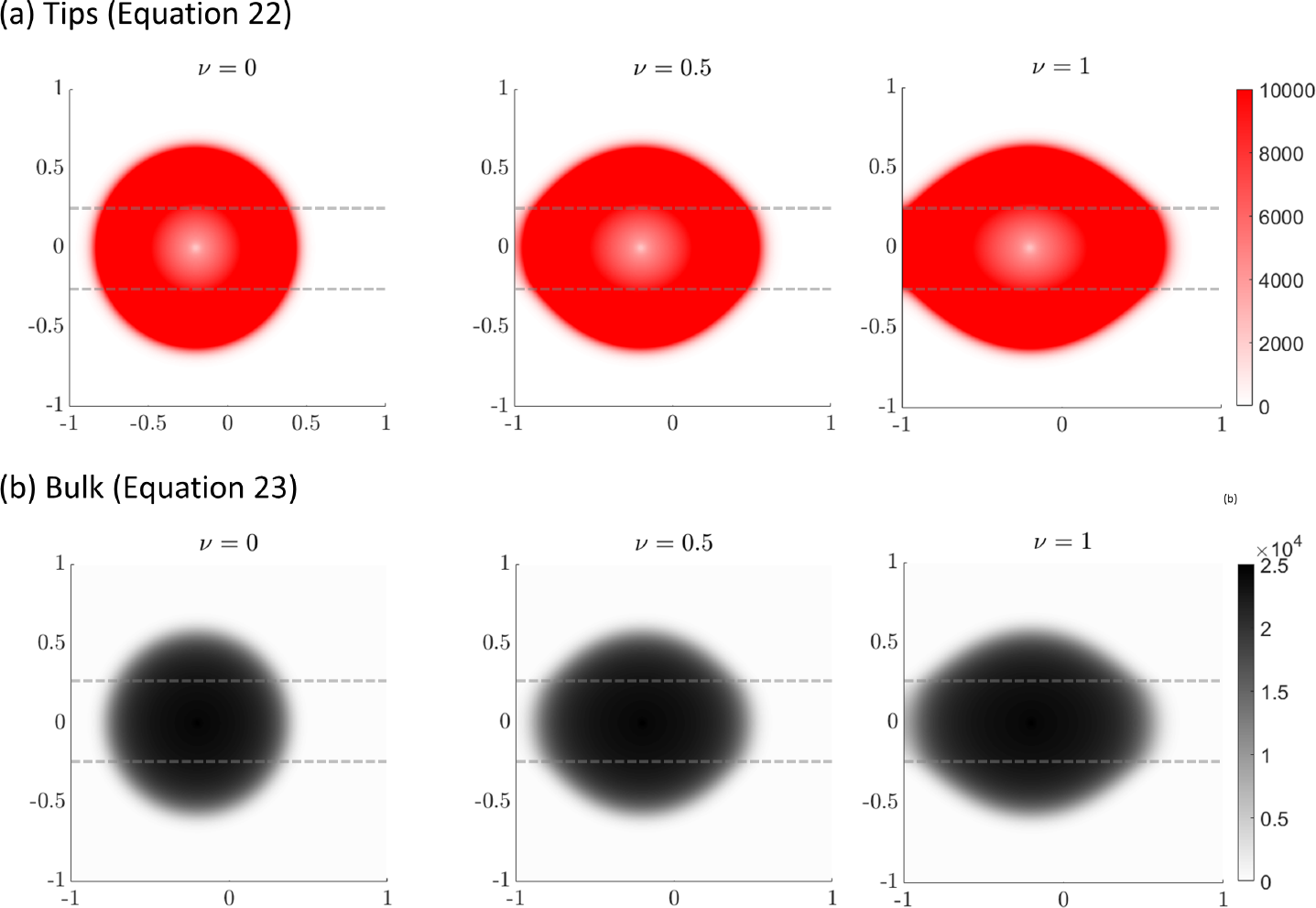
Anisotropic scenario. 2D simulations for the stripe anisotropic arrangement under different values of *ν* = 0 (first column), *ν* = 0.5 (second column), and *ν* = 1 (third column). The region of horizontally aligned fibres is enclosed by the dashed lines shown in each panel. The first row plots the distributions for the tips (red) while the second row plots the distributions for the bulk (black) for model (22-23) at T=1000 hr.

Figure 6 (b) shows simulations of the reduced tips-bulk-anisotropic model (22-23) for the same stripe arrangement. Qualitatively similar behaviour is observed, with the tumour preferentially invading along the direction of anisotropy. However, in line with earlier one-dimensional simulations, the invasion process is significantly faster. This again highlights the potential danger of applying model reductions without direct evaluation of their quantitative impact.

In Figure 7 we use model (22-23) to investigate the impact of altering parameter *ν*, a weighting parameter that controls the degree to which persistence or anisotropic alignment influences tip orientation: *ν* = 0 describes an isotropic persistent random walk, *ν* = 0.5 gives equal weights to the anisotropic and the persistent random walk, while *ν* = 1 describes a fully anisotropic random walk. As expected, increasing *ν* from 0 to 0.5 to 1 results in greater anisotropic spread of the tumour driven by enhanced invasion along the horizontal fibres.

#### 4.2.3 Test 3: Treatment

In this section, we simulate a simple tumour treatment. First, we consider the solution of the tips-bulk-anisotropic model (22, 23) under the previous stripe arrangement simulation (Figure 6) at 1000 hours. In [44] the hypothesised minimum detection threshold for the visible part of the tumour is set to be about 0.16 of the carrying capacity. In our simulated treatment, we define the treatment region to be all numerical cells where the bulk density is larger or equal to 0.16*K*. To mimic effects that could lead to the misalignment of the treatment region, such as through breathing or imperfect patient positioning, we move the treatment region a bit off-centre in the positive *y* direction by 0.05 *mm*. In the moved treatment region we set all bulk cells to zero. In the rest of the domain we model a reduced treatment effect by randomly killing bulk cells, drawn for each cell from a uniform distribution. This procedure leaves a half-moon shaped area of partially treated tumour cells behind, as seen in the third panel of Figure 8.

**Fig. 8.**
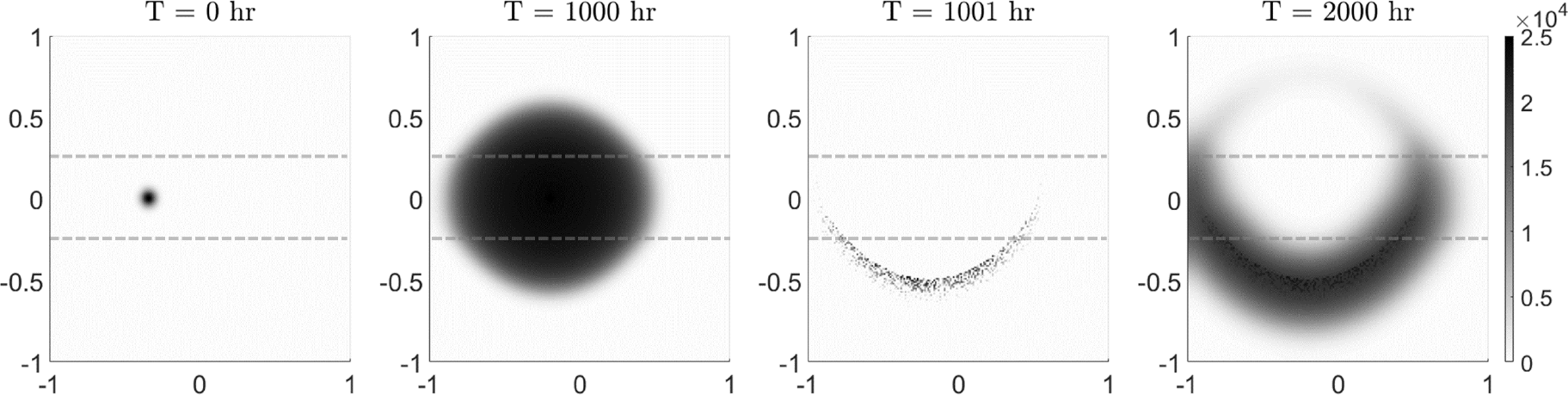
Treatment scenario. The left two panels show growth of the pre treated tumour (showing the bulk) over 1000 hours, as determined by the tips-bulk-anisotropic model (22-23) with *ν* = 1*/*2. The region of horizontally aligned fibres is enclosed by the dashed lines shown in each panel. The middle-right panel shows the bulk distribution following treatment (see text for details) while the right-most panel plots the bulk 1000 hours post treatment.

Simulations indicate that recurrence will occur even if only a small amount of tumour cells are missed during treatment. The bulk regrows from the treatment boundary as quickly as the original tumour.

## 5 Discussion

The TMT-guided glioma invasion observed by Osswald et al. [29] provides a potential paradigm shift in our understanding of glioma growth, in particular for the large class of TMT-generating astrocytomas. Video observations (see Figure 1) provide an intuitive explanation for the diffuse appearance of many gliomas. Rather than a distinct tumour boundary, a complex invasion front is formed from the interaction between TMT, moving nuclei and growing bulk cells. This invasion mechanism uncovers new treatment targets, such as the TMT or a critical TMT growth factor Cx43 [29]. Inspired by these observations, we have employed a mesoscopic transport equation framework to model the phenomenon of TMT invasion. Our model assumes that tumour growth is led through a network of thin TMT protrusions, facilitating transportation of nuclei at the invasion front which then mature into bulk cancer cells. Notably, our model assumes that transport of glioma cells is exclusively through the network-based shuttling: bulk glioma cells do not migrate independently. The production terms in our model are based on a logistic term. This could, of course, be replaced with other suitable growth laws, for example a Gompertz law.

The full model utilises kinetic transport equations within two of its equations. The analysis of such systems is far from trivial [17] and fully formulating this model required extensions to existing theory: the definition of integral operators on appropriate function spaces, identifying null spaces of operators and applying Fredholm alternatives for compact operators. While this theory is fundamental for the understanding of the scalings, it distracts from the main message of this paper (the hierarchical structure of existing and new glioma models) and the details are therefore presented in Appendix A.

Starting from the mesoscopic system of equations, we were able to find a pathway through well established macroscopic glioma models by applying a sequence of scaling limits and simplifications. In particular, this ended at the well known Fisher-KPP type model (27), commonly known as the proliferation-infiltration model (PI-model) in glioma studies [43, 44, 36]. Within our framework, a PI model can be derived when we assume that nuclei dynamics are in quasiequilibrium (A.1), that the tips-TMT dynamics is in quasiequilibrium (A.2), when we consider macroscopic time and space scales (A.3-A.4), if the spread is isotropic (A.5), and when the tips are proportional to the bulk (A.6). Consequently, a simple and tractable macroscopic model with a diffusive structure can be justified even within the context of a more complex microscopic description in which any glioma cell transport is strictly linked to the dynamics of its associated TMT network. Further, this reduction to a Fisher-KPP model allows us to find an explicit form of the invasion speed in terms of the parameters that govern the underlying glioma-TMT dynamics.

If anisotropy becomes important, for example through TMT orientation along white matter fibre tracks, then we relax (A.5) and can consider the anisotropic model (28) as done in [42, 23, 11]. If we have a clear separation of invading and growing sub-populations, we can use a go-or-grow type formulation (22,23), i.e. a structure similar to previous models in [1, 40, 15]. Typical go-or-grow models feature two different sub-populations which switch phenotype between proliferator and invader states. For the go- or-grow version here, rather, it is the TMT tips that represent the invading population.

Numerical simulations were used to compare the model types. Here we noticed that the qualitative shapes of solutions to the full model, the *tips-bulk-transport* model, and the *tips-bulk-anisotropic* model remain similar within the anisotropic diffusion tests. Therefore, the quasi-equilibrium assumptions (A.1, A.2) and the diffusive scaling (A.3, A.4) qualitatively preserve the shape of the solutions. However, the wave speed increases significantly. This is a direct result of quasi-equilibrium assumptions, which act to accelerate the underlying dynamics. Thus, ad hoc simplifications of complicated models in the interest of deriving a simpler system should be considered with care.

Modelling inevitably depends on the available data. The assumptions and parameters for our full model (1-5) is founded on detailed data obtained for a murine model of glioma growth, but similar data is not available for glioma growth in humans. Clinically, measurements and data typically rely on macroscopic imaging, such as through MRI or DTI imaging and PET imaging modalities and macroscopic models (such as the PI model) are fitted directly to such datasets, without directly connecting to the underlying microscopic processes. Here we have added to a growing literature of studies where glioma growth models are built upwards from the microscopic level, therefore providing a connection between microscopic and macroscopic parameters.

Given the complexity of the biological process, various simplifications were necessary during the modelling in order to derive a tractable system. Future investigations will involve revisiting some of these simplifications and investigating the extent to which they influence the dynamical behaviour

## Acknowledgements

NL is member of INdAM-GNFM and acknowledges the computational resources of the Department of Mathematical Sciences, G. L. Lagrange, Politecnico di Torino. KJP is a member of INdAM-GNFM.

## Data availability

All data used are from the publication of Osswald et al. [29], and are available through the original publication.

## A Structure of the Turning Operators

The model (1-5) involves kinetic transport terms in the equations of *P* and *N* . The analysis of such transport equations is involved, and it requires a carefully laid out functional analytical approach. In this section we provide these details. Comprehensive descriptions of these procedures can be found in [32, 17].

The key component of the transport process is the turning operator ℒin (1). In the following we define ℒ, consider its nullspace, find the corresponding Maxwellian *Ψ* (*v*), and derive a pseudo-inverse of ℒ.

For this let us consider the transport part of the tip equation (1)

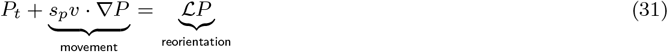

where ℒ denotes the turning operator. The turning operator acts on the *v*-dependence of *P*, hence we consider ℒ :

*L*^2^(*V*) *→ L*^2^(*V*), and we focus on the *v*-dependence in this subsection. Then for *P ∈ L*^2^(*V*) we have

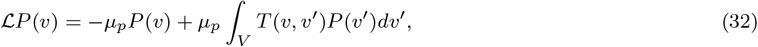

where *µ*_*p*_ is the turning rate and the turning kernel is *T* (*v, v*^*i*^). We note that microtubes follow the direction of nerve fibres, but also keep relatively straight and do not bend too much. To model both effects, we use a combination of a trail following term ℒ_1_ and a persistent random walk ℒ_2_ such that ℒ is a convex combination

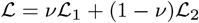

for some 0*≤ ν≤* 1. The directed component ℒ_1_ has the kernel *T*_1_(*v, v*^*i*^) = *q*(*v*), where *q*(*v*) is a probability distribution representing the distribution of nerve fibre directions. Note that in the full model *q*(*x, v*) will depend on space. Here we focus on the *v*-dependence only. Then

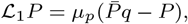

where 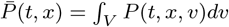 A persistent random walk can be modelled using a cosine bias of the previous direction [32] as

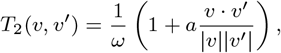

for a persistence parameter 0 *< a <* 1 and *ω* = |*V* |. Then

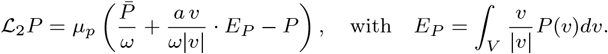

Through this construction we can also identify a full turning kernel as the convex combination of *T*_1_ and *T*_2_.

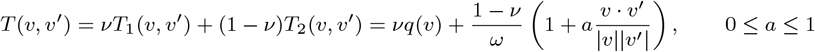

where *ν ∈* [0, 1]. Let us suppose that *q* is bidirected, so that

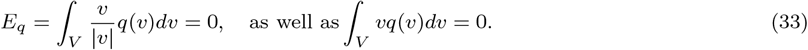

If we consider scaling limits, then the null space of the operator ℒ becomes important. Each of the operators ℒ_1_ and ℒ_2_ have been studied in isolation many times ([32, 17]). We have that ker ℒ_1_ = *q*, while ker ℒ_2_ = 1 . As we show next, the kernel of ℒ is a linear combination of these:

### Lemma 1

*We consider L*^2^(*V*) *with the weighted inner product*

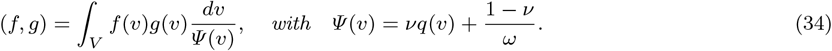

*We assume*

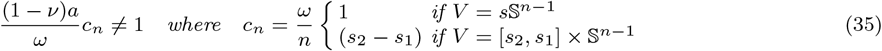

*Then*

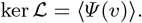

Note that in the above Lemma we ignored the possible *x* dependence in *q*. If *q* depends on *x* then *Ψ* does as well. The function *Ψ* (*v*) is a probability density function as

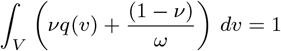

is called the *equilibrium distribution* with ℒ *Ψ* = 0.

**Proof**. We first show that *Ψ ∈* ker ℒ. Doing so requires showing that ℒ *Ψ* = 0. Written in explicit terms, this is

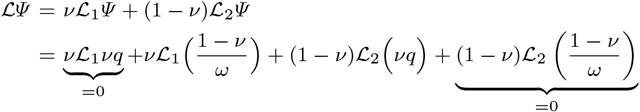

where the two 0-terms follow from direct calculation. The mixed terms demand special attention

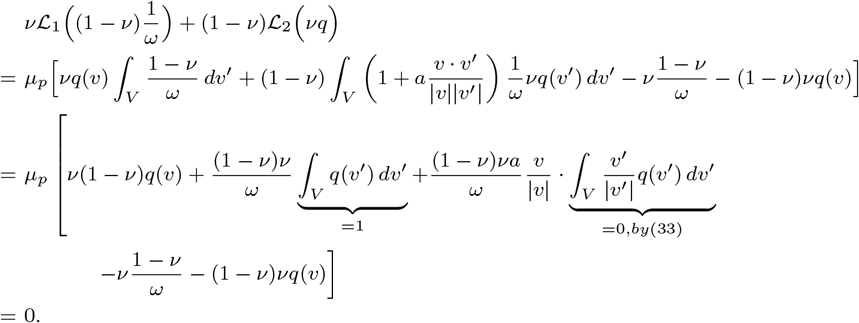

Let us now prove that ker ℒ *⊂ Ψ* . Consider *ϕ ∈* ker ℒ and, for the sake of argument, suppose that *ϕ* does not belong to *Ψ* . Then, *ϕ* has a component in *Ψ* ^*⊥*^, which we again call *ϕ* for simplicity. Then

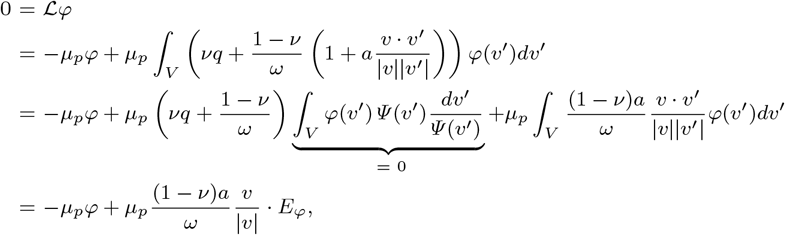

where

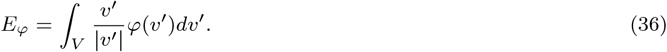

This implies that *ϕ* has a specific form

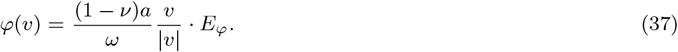

It is straightforward to check that this expression is indeed perpendicular to *Ψ* (*v*):

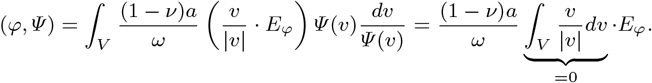

We substitute (37) into (36) to obtain

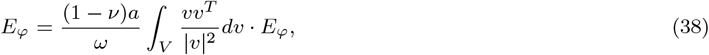

which is a condition on *E*_*ϕ*_. To further analyze this condition, we need to compute the second velocity moment

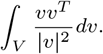

Such calculations have been performed in [16] and we summarize the important steps here. We distinguish two cases. In the first case we assume a constant particle speed, then the set of velocities becomes 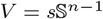. Using the substitution *v* = *sθ* with unit vector *θ*, we find

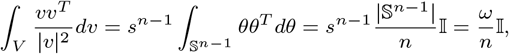

where 𝕀 is the identity matrix and *ω* = |*V* |.

In the second case we consider *V* = [*s*_1_, *s*_2_] *×*𝕊 _*n−*1_, where particles can have speeds in the interval [*s*_1_, *s*_2_]. Then

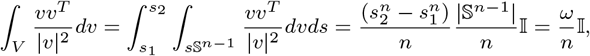

where again *ω* = |*V* |. Then condition (38) becomes

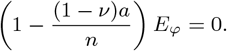

Since 0 *< ν <* 1, *a <* 1 and *n >* 1, we have that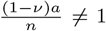 and this implies that *E*_*ϕ*_ = 0. Then, from (37), we also have *ϕ* = 0 and consequently *ϕ ∈ Ψ* .

QED.

